# Autophagy is required for midbrain dopaminergic axon development and their responsiveness to guidance cues

**DOI:** 10.1101/2020.10.09.333435

**Authors:** M.S. Profes, A. Saghatelyan, M. Lévesque

## Abstract

Mesodiencephalic dopamine (mDA) neurons play a wide range of brain functions. Distinct subtypes of mDA neurons regulate these functions but the molecular mechanisms that drive the mDA circuit formation are largely unknown. Here we show that autophagy, the main recycling cellular pathway, is present in the growth cones of developing mDA neurons and its level changes dynamically in response to guidance cues. To characterize the role of autophagy in mDA axon growth/guidance, we knocked-out (KO) essential autophagy genes (*Atg12, Atg5)* in mice mDA neurons. Autophagy deficient mDA axons exhibit axonal swellings and decreased branching both *in vitro* and *in vivo*, likely due to aberrant microtubule looping. Strikingly, deletion of autophagy-related genes, blunted completely the response of mDA neurons to chemo-repulsive and chemo-attractive guidance cues. Our data demonstrate that autophagy plays a central role in regulating mDA neurons development, orchestrating axonal growth and guidance.

## Introduction

During mammalian embryogenesis, neurons form a highly interconnected network by elongating and projecting their processes toward their target areas. Axonal development is organized into multiple steps starting from growth cone formation, axonal growth and branching, which are coordinated by both intrinsic mechanisms and extrinsic cues from the surrounding microenvironment that shape axonal growth and guidance (Tamariz and Varela-Echavarría, 2015). Midbrain dopaminergic (mDA) neurons located in the ventral tegmental area (VTA) and *substantia nigra pars compacta* (SNpc) project their axons to different brain regions and control a wide range of behavioral responses. VTA neurons are medially located in the ventral midbrain and project to different regions implicated in the reward pathway, including the *nucleus accumbens* and the prefrontal cortex (PFC), constituting the mesocorticolimbic pathway. Imbalances of dopamine input in the mesocorticolimbic pathway have been linked to depression, drug addiction, attention deficit hyperactivity disorder and schizophrenia (Van den Heuvel Pasterkamp, 2008). Laterally located SNpc neurons target the dorsolateral striatum (caudate and putamen), forming the nigrostriatal pathway. Nigrostriatal connectivity loss is the main histopathological feature of Parkinson’s disease (PD) (Van den Heuvel and Pasterkamp, 2008). Several axonal guidance molecules have been described and shown to be crucial for the proper growth and targeting of mDA neuron axons. Sema7a and Netrin-1 seem to be of special importance in guiding such axons to their proper targets in the striatum and in the PFC (Chabrat et al., 2017; Flores, 2011).

Despite the growing body of evidence that describes axonal guidance and guidance molecules and their receptors, the downstream effectors of such important proteins remain elusive. Interestingly, mRNA translation, Ca^2+^ signaling and local protein degradation have been implicated as downstream effectors of axonal guidance (Campbell and Holt, 2001; Sutherland et al., 2014). Additionally, endocytosis and exocytosis in the growth cone have been suggested to be involved in the specific turning of the growth cone toward or away from certain environmental cues (Tojima et al., 2011). Interestingly, there is commonly cross-talk between the endocytosis and autophagy pathways, resulting in endocytic cargo being recycled by autophagy (Lamb et al., 2013). Furthermore, macroautophagy is the major catabolic mechanism in eukaryotic cells that is responsible for the removal of long-lived proteins and damaged organelles in the lysosome (Kaur and Debnath, 2015; Lee et al., 2012). This dynamic self-digesting process will hereafter be referred to as autophagy. Autophagy is a very conserved process and relies on several protein complexes that are well characterized and are implicated from the initiation to the degradation step (Kaur and Debnath, 2015; Lee et al., 2012). Autophagy proteins have been identified and named autophagy-related proteins (ATGs) (Kaur and Debnath, 2015; Lee et al., 2012). In virtually all cells, autophagy is active at a basal level, performing homeostatic functions such as protein and organelle turnover. This housekeeping process is upregulated in response to both extracellular and intracellular stress conditions, such as nutrient starvation, growth factor withdrawal, hypoxia, aggregation of proteins, accumulation of damaged mitochondria, oxidative stress and high bioenergetic demands (Kaur and Debnath, 2015; Lee et al., 2012). Autophagy is also stimulated when cells undergo structural remodeling (Mizushima and Levine, 2010). Notably, interfering with the expression of different autophagy proteins has been reported to cause brain defects. Knockout embryos for Ambra1, an autophagy-related protein, display accentuated mid-hindbrain exencephaly (Fimia et al., 2007), whereas Atg7 conditional knockout (cKO) in mDA neurons leads to aberrant axonal morphology and altered dopamine release in young animals (Hernandez et al., 2012). Moreover, Alfy/WDFY3, an autophagy adaptor protein, has been shown to regulate the formation of major axonal tracts in the brain as well as in the spinal cord (Bosco et al., 2016), and microRNA Mir505-3p, which targets *Atg12*, a core autophagic protein, regulates axonal elongation and branching *in vitro* and *in vivo* (Yang et al., 2017). Much of this evidence, however, comes from experiments with autophagic adaptor and/or regulatory proteins, not core autophagic proteins; therefore, it cannot be excluded that some of the brain defects reported might be related to their non-autophagic roles. Additionally, most of the KO models used were nonneuronal-specific; thus, one cannot exclude the possibility that the observed axon defects could be due to alterations in the cellular organization of nonneuronal cells in the brain. Finally, little is known about the role of autophagy in axonal growth and the guidance of dopaminergic neurons and how this major catabolic pathway is regulated in response to different microenvironmental cues. Therefore, more data are needed to fully corroborate a central role of autophagy in brain development and its implications in the axon growth/guidance of dopaminergic neurons.

In this study, using CRISPR-Cas9 gene editing and cKO of Atg5 specifically in dopaminergic neurons, we show that autophagy is required for proper mDA axonal morphology and branching during development. Moreover, we demonstrate that autophagy plays a central role in mediating the responses of dopaminergic axons to guidance cues, and ablation of Atg5 in mDA neurons completely blunts axon growth/guidance in response to major chemorepellant and chemoattractive signals, such as Sema7a and Netrin-1. Our data reveal a central role of autophagy in dopaminergic system development by regulating axonal growth/guidance and mediating the responses of these cells to extrinsic guidance cues.

## Results

### Autophagy machinery is present in mDA neuronal soma and axons and is enriched in growth cones

To determine whether autophagy could potentially be involved in mDA development, we first performed immunofluorescent staining on brain sections at different maturational stages. The presence of autophagy was assessed in midbrain sections collected from embryonic day 14 (E14) to postnatal day 7 (P7) through TH (tyrosine hydroxylase) marker and LC3 (an essential autophagy protein) colabeling. This developmental window from E14 to P7 corresponds to the entry of mDA axons in the ganglionic eminence (E14.5) and was chosen because mDA axons are growing toward their forebrain targets in the embryonic stage, having their connections consolidated at approximately P7. LC3 is expressed in TH+ mDA neurons in the VTA and their axon projections in the medial forebrain bundle throughout development (**Fig. 1A and B**). To investigate the expression of LC3 in growth cones (GCs), we employed explant cultures. Immunofluorescent signals for LC3 were highly expressed in TH+ GCs from 3 day *in vitro* (3DIV) explant cultures (**Fig. 1C**). To further characterize autophagy in mDA development, we performed western blotting from ventral midbrain samples collected from wild-type mice. Samples were collected from embryonic stages to adulthood to comprise the developmental window described above. Our western blot analysis of the autophagic marker LC3-II, which represents the lipidated autophagosomal-associated form of LC3; the ATG5-ATG12 complex, which is required for autophagy initiation; and p62, which is the main autophagic substrate, showed similar temporal profiles. These markers displayed increasing levels from embryonic to postnatal stages, reaching their highest levels at P7 with a subsequent decrease with further postnatal development (**Fig. 1D**). Interestingly, P7 corresponds to a developmental stage where axonal pruning occurs, a mechanism that requires dynamic axonal changes (Hu et al., 2004). This structural plasticity is required for mDA connection consolidation and fine-tuning. Hence, the presence and temporal regulation of autophagy markers throughout mDA neuron development and LC3 enrichment in GCs suggest that autophagy may play a major role in dopaminergic circuit formation and axon growth/guidance.

**Figure 1:**
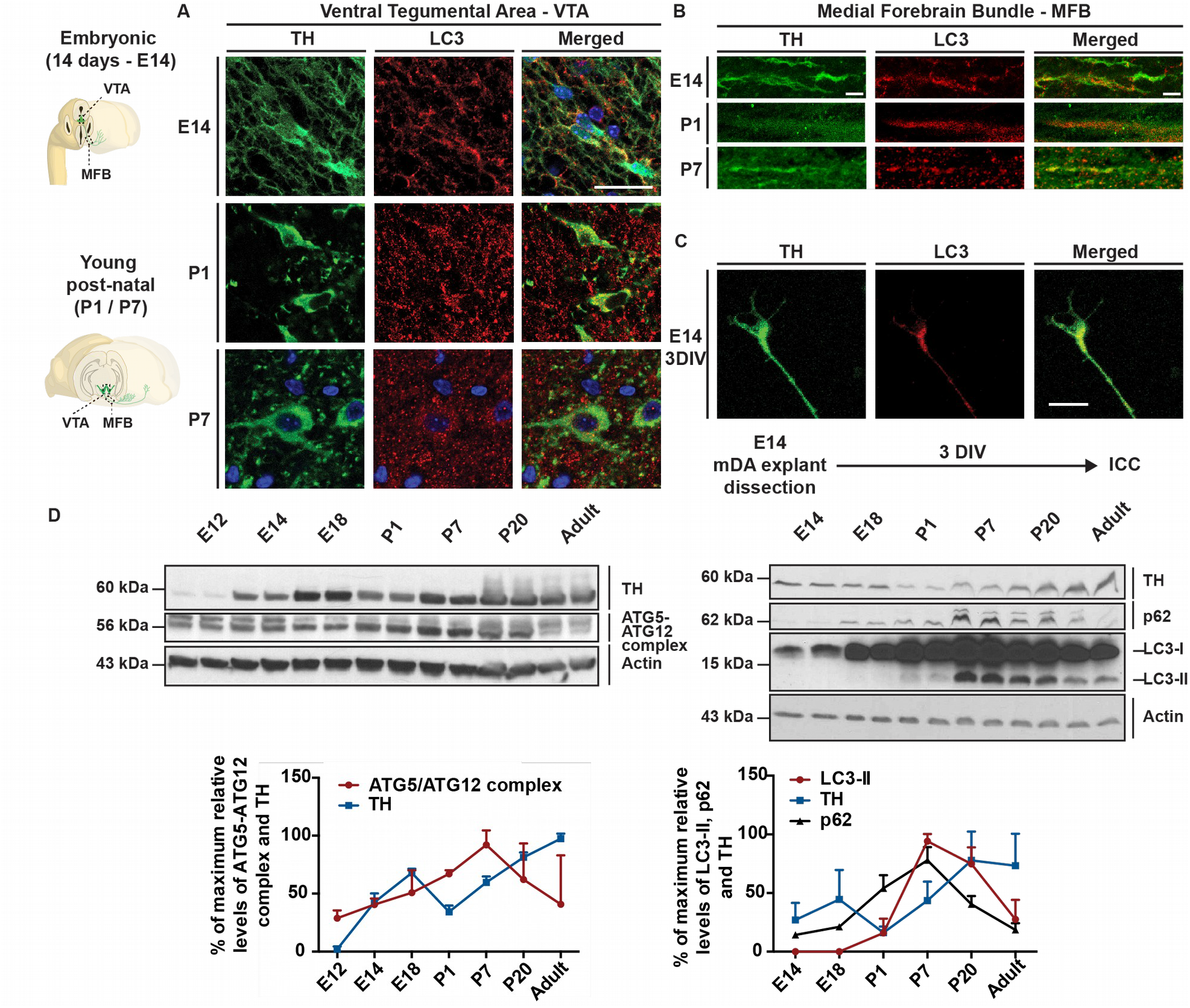
Autophagy is present and is timely regulated in midbrain dopaminergic neurons and axons throughout development. Representative confocal images of immunohistochemical analysis of the autophagic marker LC3 in the VTA (A) and in the medial forebrain bundle (B) from E14 to P7 (n=3 animals with an average of 3 sections per animal). (C) Immunolabeling for LC3 in E14 explant cultures showing enrichment of autophagy markers in the GCs of TH+ axons (n=3 explants from 3 individual embryos). (D) Western blot analysis of autophagic markers LC3-II, ATG5-ATG12 complex and p62 displays time-dependent changes in the protein levels during the developmental window (embryonic to postnatal stages, n=3 animals per age). Scale bars: A, 25 μm, and B and C, 10 μm.

### Autophagy regulates mDA axonal morphology and arborization

To test the role of autophagy during mDA circuit development, we generated autophagy-deficient conditional mutant mice in mDA neurons by crossing *Dat*^*Cre*/+^ mice with *Atg5*^*flox/flox*^ mice (hereafter referred to as ATG5 cKO). Dat^+/+^Atg5f^*/f*^ mice were used as controls, unless noted otherwise in the text. *Cre* recombinase expressed under the control of the *Dat* promoter was efficient in deleting *ATG5* from mDA neurons as seen by western blotting performed on P7 ventral midbrain samples (Fig. S1). Importantly, because of the DAT expression pattern, Atg5 deletion occurs only in postmitotic mDA neurons just prior to their axonal growth toward the forebrain (Prestoz et al., 2012), allowing blockage of autophagy at a developmental time point that does not interfere with mDA neuron differentiation from their progenitors.

Since autophagy is known to be involved in cytoskeletal remodeling, we first examined its requirement for mDA axonal growth and morphology. We used explants of the VTA of E14 ATG5 cKO mice and their control littermates cultured for 3DIV and measured axon diameter and GC area. Axons and GCs from ATG5 cKO explants displayed an abnormal morphology and showed 2.6- and 1.7-fold increases in the axon diameter and GC area, respectively, compared to axons in the control explants (**Fig. 2A**). Sholl analysis also showed a reduced number of intersections of axons in ATG5 cKO explants compared to control explants (**Fig. 2B**). Importantly, the length of axons and their number that initially grew outward from the explants did not differ significantly between mutants and control explants. To test whether similar morphological defects are observed following manipulation of other autophagy-related genes in mDA neurons, we next performed CRISPR-Cas9 gene editing for *Atg12,* a core autophagy-related protein required for autophagic vesicle formation (Kaur and Debnath, 2015; Lee et al., 2012). Similar to ATG5 cKO, CRISPR-Cas9 gene editing for *Atg12* led to GCs and axonal enlargements with similar fold-change to ATG5 cKO (2.6- and 1.3-fold changes, respectively; Fig. S2).

**Figure 2:**
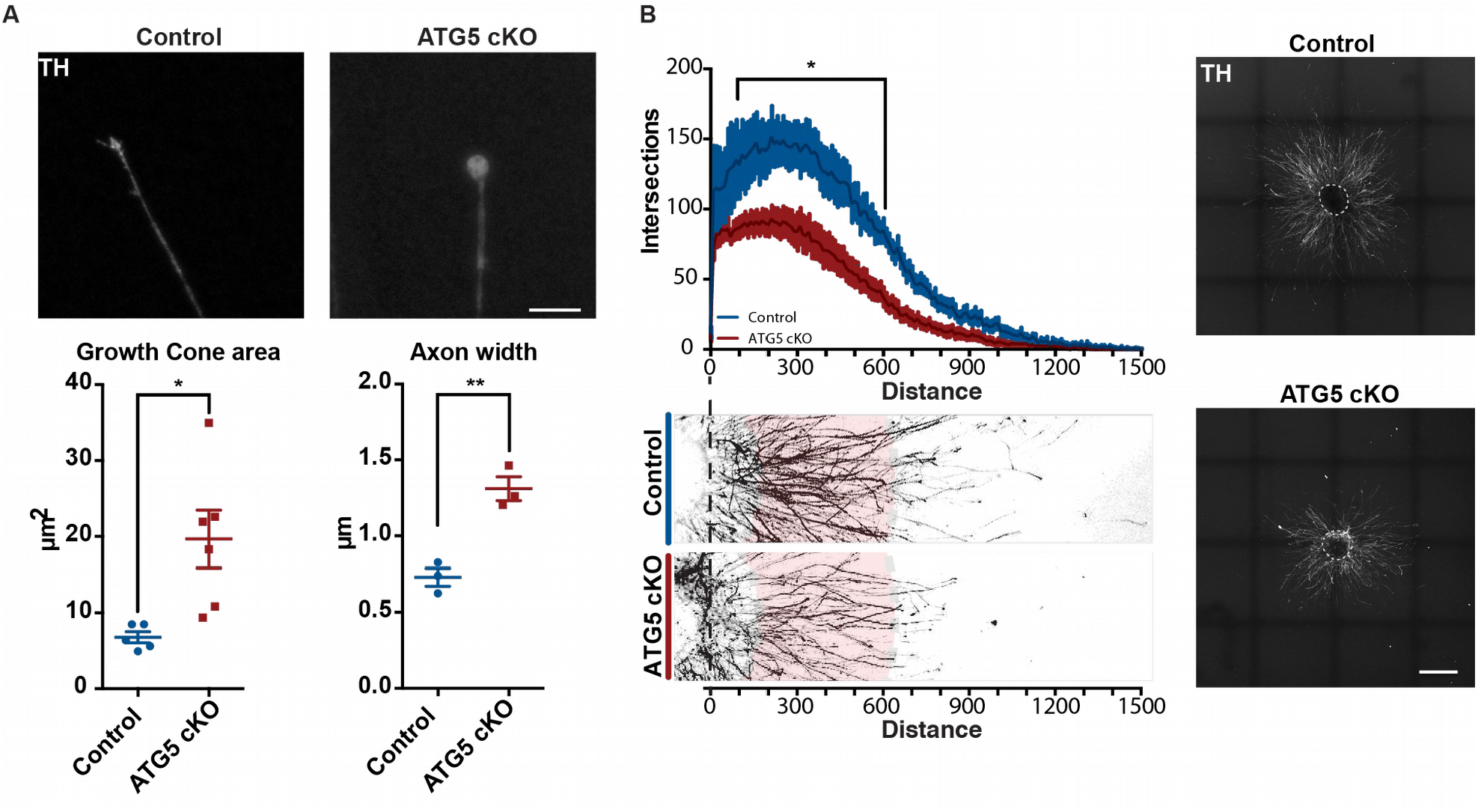
Autophagy ablation leads to morphological changes in mDA axons and GCs. (A) E14 VTA from (a) *Dat*^+/+^*Atg5*^*flox/flox*^ (control) and *Dat*^*Cre*/+^*Atg5*^*flox/flox*^ (ATG5 cKO) mice were dissected into explants and cultured in 3D collagen matrix for 3DIV prior to fixation and ICC. GC area: n=5 explants for control samples and n=6 explants for ATG5 cKO samples dissected from 4 animals for each genotype. For each explant, an average of 20 GCs were analyzed, and the average per explant is plotted (mean values= 6.78 for control and 19.66 for ATG5 cKO; *p*=0.0142; two-tailed t test). Axon width: n=3 explants dissected from 3 animals for each genotype per condition with an average of 20 axons analyzed per explant and averaged (mean values= 0.73 for control and 1.3 for ATG5 cKO; *p*= 0.0041; two-tailed t test). (B) Sholl analysis of E14 explants at 3DIV reveals axonal arborization differences between ATG5 cKO and wild-type samples. n=7 explants for controls and n=10 for ATG5 cKO samples dissected from an average of 6 animals. Two-way ANOVA and the Sidak test were used for post hoc comparisons, *p < 0.05). Scale bar A, 10 μm, and B, 500 μm.

To determine the requirement of autophagy for mDA axonal morphology *in vivo,* we performed immunohistochemistry for TH on P11 nigrostriatal brain sections derived from ATG5 cKO and control mice. P11 was chosen since the nigrostriatal dopaminergic connections are established around P7, which allows us to assess the impact of autophagy on the establishment and/or maintenance of mDA axons. To ensure that the lack of autophagy ablation did not induce the loss of dopaminergic neurons and that any possible defects in the number or density of axonal profiles in the striatum were not due to lower number of mDA neurons, we performed stereological counting of TH+ neurons in the SNpc and VTA. Neuronal counting did not reveal any difference in the total number of mDA neurons between mutant and control mice (**Fig. 3A and B**). To study the axonal projection morphology *in vivo*, we counted the number and size of TH+ profiles (see arrowheads in **Fig. 3A**) on matched coronal sections from ATG5 cKO mice and control littermates. Our analysis revealed a 1.5-fold enlargement of TH+ axonal profiles and a 0.7-fold reduction in their number when comparing mutants and controls. These data are in line with our *in vitro* analysis showing enlarged GCs and reduced axonal branching and indicate that autophagy is required for proper morphological development and/or maintenance of mDA axons.

**Figure 3:**
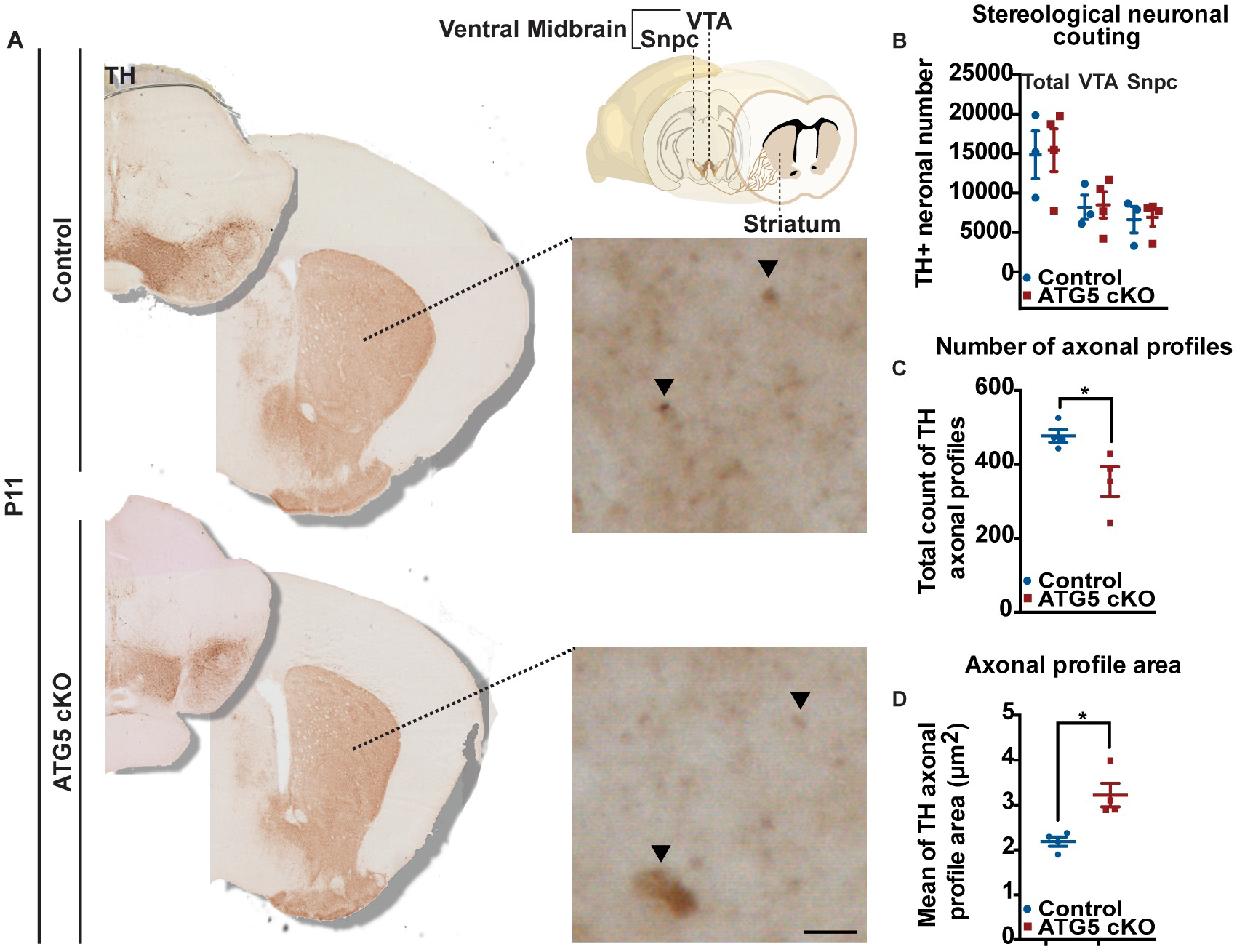
Autophagy regulates dopaminergic axonal morphology and arborization *in vivo*. (A) Immunohistological images reveal different striatal TH staining patterns when comparing P11 controls and ATG5 cKO mice. A higher-magnification image shows the abnormal axon coronal profile enlargements of TH+ axons. (B) mDA neuronal stereological counting shows no difference between groups, n=3 animals per condition (3 sections per animal). Quantification of higher-magnification images showing a reduced number of coronal axon profiles reaching the striatum (C) and the enlargements of the aforementioned profiles (D). N=4 animals per condition (3 sections per animal) and *p*=0.03 (mean values= 477.23 for control and 353.27 for ATG5 cKO) and *p*=0.0104 (mean values= 2.18 for control and 3.22 for ATG5 cKO) for C and D, respectively (two-tailed t test). Scale bar, low magnification 50 μm and higher magnification 10 μm.

### Autophagy ablation leads to the formation of aberrant microtubule loops within GCs

It has been previously shown that enlarged GCs can be characterized by a round-shaped tip with looped microtubules (Dent et al., 1999). We thus tested whether autophagy ablation could lead to the aberrant formation of microtubule loops. We performed primary ventral midbrain culture from ATG5 cKO mice and control littermates and performed immunostaining for TH as an mDA neuronal marker in combination with alpha-tubulin immunolabeling to visualize microtubule organization. GCs from midbrain cultures of ATG5 cKO mice displayed an increased prevalence of intra-axonal loops compared to controls (**Fig. 4**). Measurement of these loops also revealed that microtubule loops from ATG5 cKO mice were 1.7-fold larger than the loops found in the GCs of control animals. TH+ GCs containing two or more microtubule loops were also observed, and the percentage of GCs with 2 or more loops was much more frequent in ATG5 cKO cultures than in control cultures (2.3-fold change). In sum, autophagy ablation leads to abnormal microtubule phenotypes in dopaminergic GCs and most likely underlies the enlarged GCs observed in ATG5 cKO Gcs.

**Figure 4:**
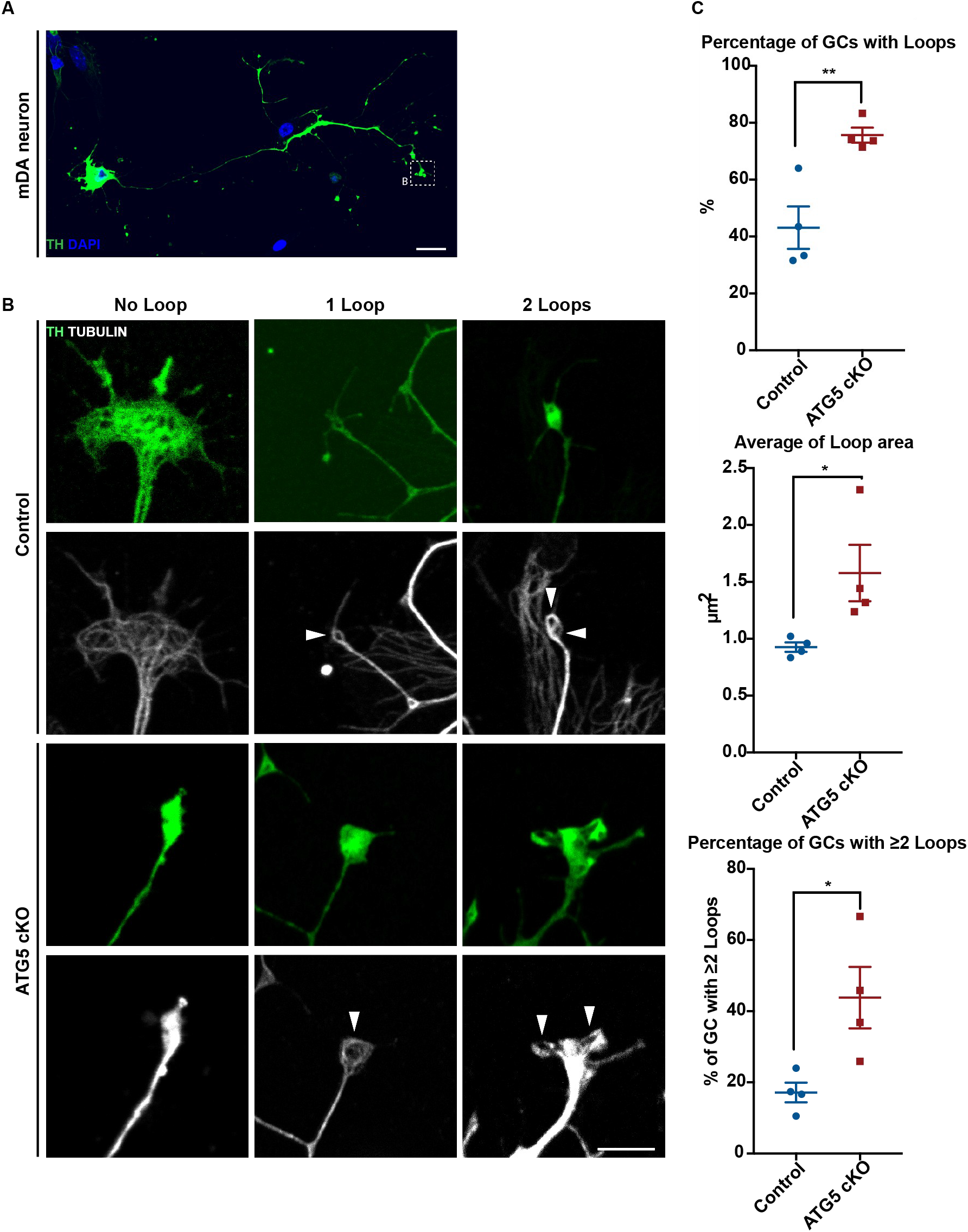
Autophagy ablation in mDA neurons leads to the formation of aberrant microtubule loops in GCs. (A) Representative immunohistological image of an mDA primary culture neuron at 3DIV. Dashed B square depicts an example of GC. (B) Representative high-magnification images of GCs colabeled with TH and tubulin. From left to right, images are separated into GCs without any loop, those with one loop and those with two loops. (C) Quantification of high-magnification images showing from top to bottom that ATG5 cKO cultures display a higher prevalence of loops in GCs than controls (75% in comparison to 43%; *p*=0.0062), a 1.7-fold larger area (control mean value= 0.92; ATG5 cKO mean value= 1.57; *p*=0.0413) and a higher prevalence of 2 or more loops within GCs (control= 17%; ATG5 cKO= 43.8%; *p*=0.0259). Arrowheads indicate loop presence in the image. The top two rows are images derived from control cultures, and the bottom rows are images from ATG5 cKO cultures. The results shown here are averaged from 4 independent experiments (n=4) with an average of 10 GCs analyzed per experiment. Scale bar, low magnification 50 μm and higher magnification 10 μm.

### Autophagy is induced in GCs upon guidance cue exposure in mDA primary cultures

Autophagy has been shown to regulate the cytoskeleton (He et al., 2016), cellular energy balance (Kaur and Debnath, 2015) and receptor turnover (Khan et al., 2014), all of which have been reported to be involved in axon guidance. Hence, we hypothesized that autophagy could regulate GC responsiveness to guidance cues. To test this hypothesis, wild-type mDA primary cultures were incubated with Sema7a (250 ng/mL), Netrin-1 (200 ng/mL) or control vehicle (1× PBS) for 4 h at 3DIV followed by immunolabeling for TH and LC3. Sema7a and Netrin-1 are important guidance cues for mDA axons, controlling proper innervation of the striatum and the prefrontal cortex. Since autophagosomal LC3 is represented by a dotted pattern (Klionsky et al., 2016), we quantified the number of LC3 puncta in TH+ axons and GCs. Cultures treated with Sema7a and Netrin-1 displayed 1.6- and 1.5-fold increases in the number of LC3 dots, respectively, in the GCs when compared to control conditions (**Fig. 5A-D)**. When analyzing the number of LC3 dots found in the axon (40 μm away from the GC), the results show a decrease in the number of LC3+ dots following Sema7a and Netrin-1 treatment (0.6- and 0.44-fold decreases, respectively, **Fig. 5C and B**). These data indicate a shift in the localization of the autophagosomes rather than changes in the overall level of autophagy after exposure to guidance cues. In line with this, western blot quantification of LC3-II protein levels from total culture extracts upon guidance cue exposure did not demonstrate any differences between control, Sema7a and Netrin-1 treatments (**Fig. 5E**). These data suggest that under our experimental conditions, guidance cue treatment induces little or no autophagosomal biogenesis. Rather, guidance cue stimuli affect the dynamics of already existing autophagosomes and induce their local recruitment to GCs.

**Figure 5:**
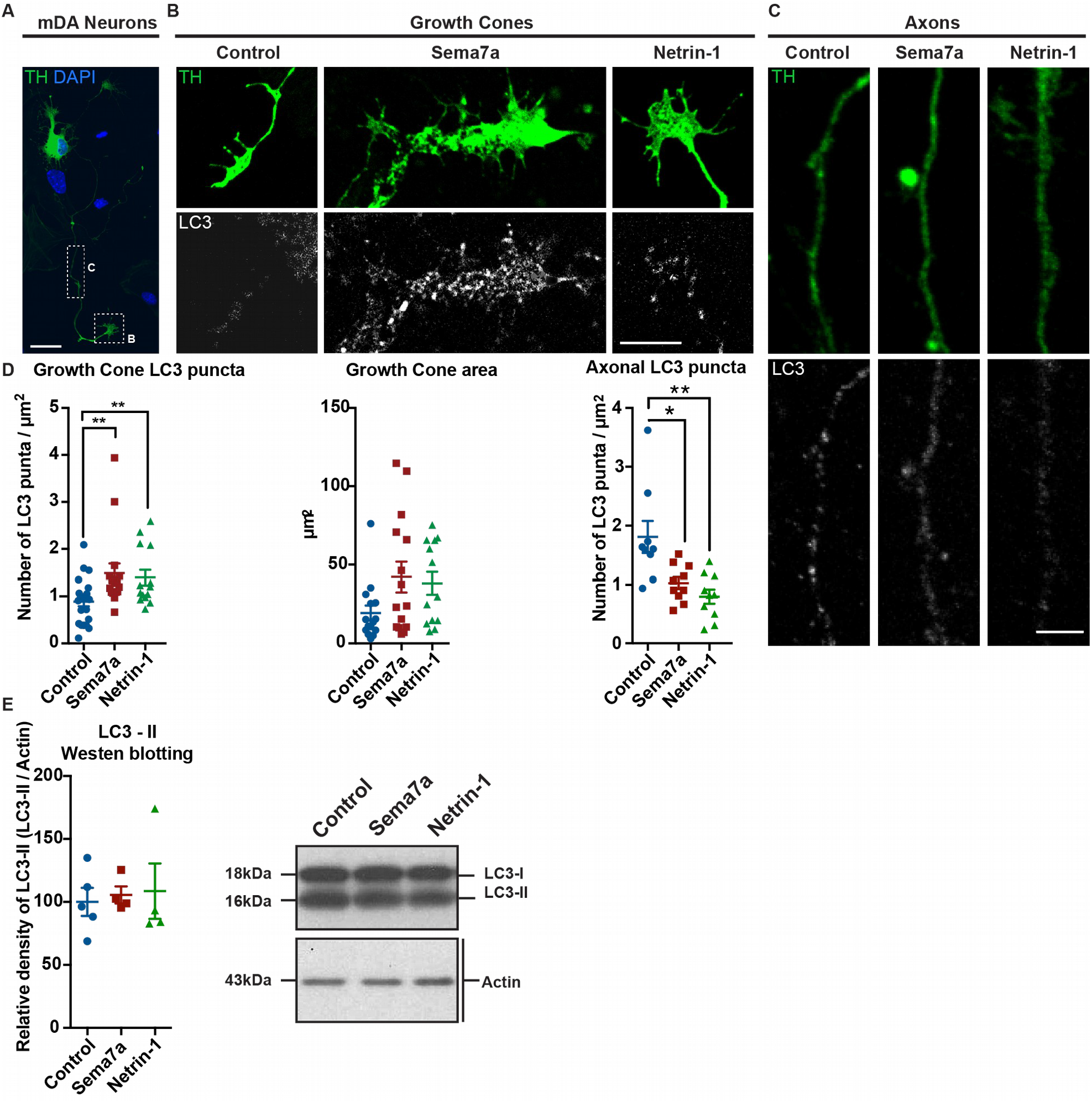
Sema7a and Netrin-1 regulate autophagy in mDA axons and GCs. (A) Representative confocal lower-magnification image of mDA 3DIV primary culture neuron. Dashed B square in the image depicts the identification of axonal GCs and dashed C rectangle depicts the axonal region quantified (40 μm away from the GC). (B) Representative higher-magnification images of TH and LC3 immunolabeling in GCs (B) and axons (C) after 4 h of treatment with PBS (control), Sema7a (250 ng/mL) and Netrin-1 (200 ng/mL). (D) LC3 dots quantification within the GCs [two-tailed t test, *p*=0.0083 for control (mean value= 0.89; n=22 GCs) vs. Sema7a (mean value= 1.48; n=14 GCs)-treated GCs and *p*=0.0096 for control vs. Netrin-1 (mean value= 1.39; n=14 GCs)-treated GCs] and axons [two-tailed t test, *p*=0.0122 for control (mean value= 1.8; n=9 axons) vs. Sema7a (mean value= 1.02; n=10 axons)-treated axons and *p*=0.0027 for control vs. Netrin-1 (mean value= 0.8; n=10 axons)-treated axons] after guidance cue exposure. GC area quantification revealed a trend to increase after cue exposure. (E) Western blot analysis of 3DIV mDA primary cultures treated for 4 h with PBS (control), Sema7a (250 ng/mL) and Netrin-1 (200 ng/mL). The results shown here are from at least 3 independent experiments.

### Autophagy ablation blunts Sema7a and Netrin-1 guidance effects in mDA cultures

Since autophagy is dynamically regulated in response to guidance cues, we next asked whether autophagy disruption could lead to altered responses of GCs to these guidance cues and affect axonal dynamics and growth. We thus performed time-lapse imaging of GCs *in vitro* and quantified the average displacement rate under baseline, Sema7a and Netrin-1 applications in control and ATG5 cKO cells. ATG5 cKO axons displayed a lower average displacement rate than axons derived from wild-type animals (**Fig. 6A and B**), underscoring the importance of autophagy in axonal growth homeostasis. Interestingly, while Sema7a application decreased the average displacement rate of control GCs, no effect of this chemorepulsive molecule on GCs of ATG5 cKO was observed (**Fig. 6A, B and C**). Similarly, upon application of the chemoattractant Netrin-1, control axons increased their average displacement rate; however, this effect was blunted in ATG5 cKO axons (**Fig. 6A, B and D**). These striking results indicate that autophagy mediates the effects of chemorepulsive and chemoattractive cues on GC growth and dynamics in mDA neurons and that a lack of autophagy completely blunts neuronal responses to guidance cues. Since Sema7a is a membrane-associated glycophosphatidylinositol-anchored (GPI-anchored) protein and not a diffusible cue, we also tested the axon response of ATG5 cKO and control mDA explants grown on alternating stripes of Sema7a. Axons from mDA explants cultured in a Sema7a stripe assay displayed a clear avoidance for Sema7a stripes when provided with a choice between the control and Sema7a substrate (**Fig. 6F**). Strikingly, axons from ATG5 cKO explants in the same conditions avoided significantly fewer Sema7a stripes (**Fig. 6F**). Altogether, our observations show that autophagy is required downstream of Sema7a and Netrin-1 signals in mDA axon guidance.

**Figure 6:**
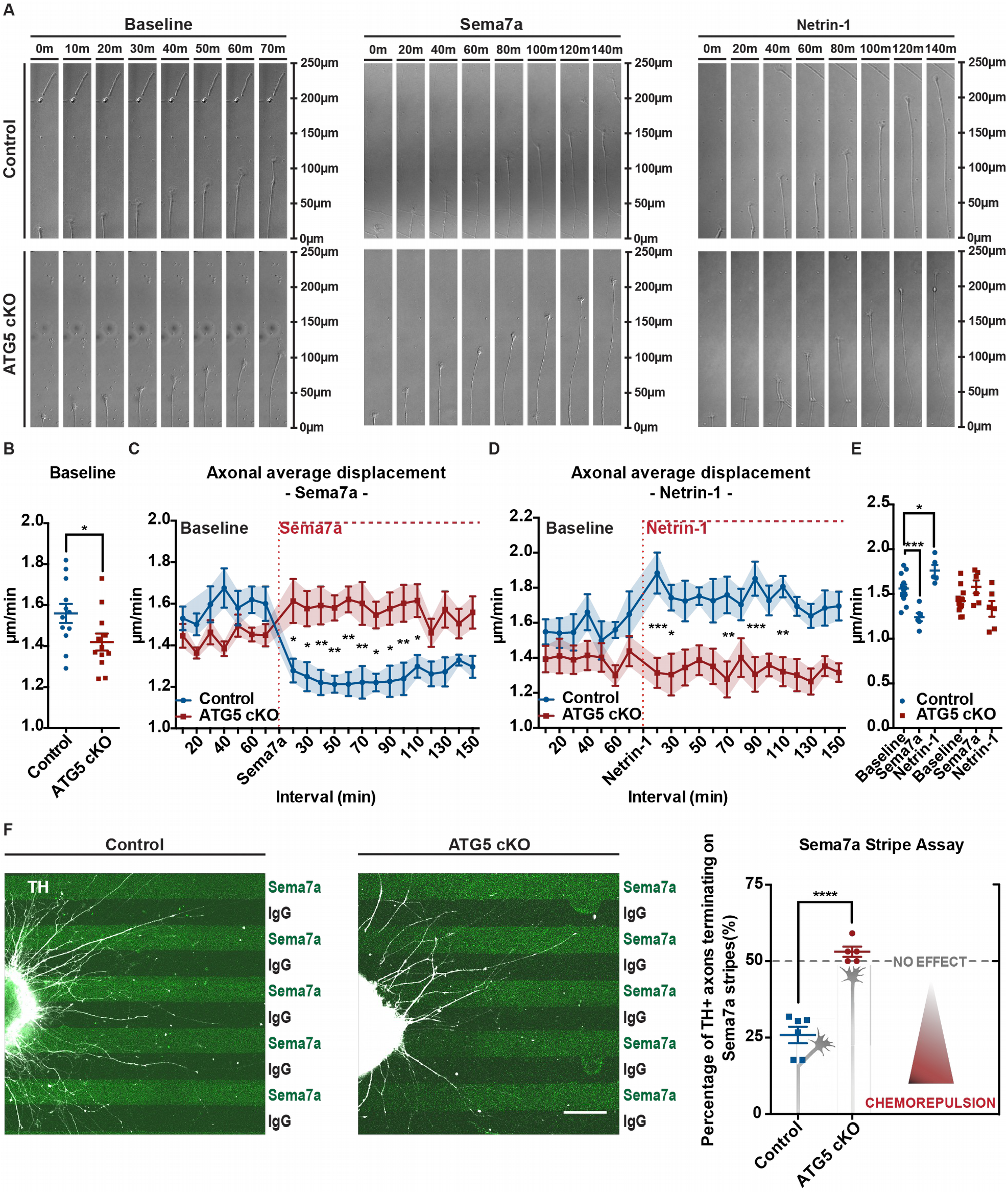
Autophagy is required for GC responsiveness to Sema7a and Netrin-1 guidance cues. (A) Time-lapse images from control and ATG5 cKO axons before and after guidance cue. (B) Baseline average displacement rate (n=12 explants per condition, average of 10 GCs per explant, two-tailed t-test; mean values=1.55 for control and 1.41 for ATG5 cKO; *p*=0.0357). (C) Displacement curves including baseline and postexposure intervals for Sema7a and (D) Netrin-1 (n=6 explants per condition with an average of 10 GCs analyzed per explant, two-way ANOVA with post hoc Sidak test, \**p*< 0.05, \*\**p* < 0.005 and \*\*\**p* < 0.0005); and (E) grouped per genotype analysis of average baseline displacement, post Sema7a and Netrin-1 exposure average displacement rate [baseline (n=12 explants, an average of 10 GCs) and Sema7a and Netrin-1 treatments (n=6 explants, an average of 10 GCs), two-tailed t-test *p*=0.0005 for control/PBS vs. control/Sema7a and *p*=0.0285 for control/PBS vs. control/Netrin-1 (mean values=1.55 for control/PBS; 1.21 for control/Sema7a and 1.76 for control/Netrin-1). (F) VTA axons stripe assay images and quantification (n=6 explants for the control and 5 for the ATG5 cKO; data are from 5 independent experiments). Two-tailed unpaired t-test; mean values=0.51 for control and 1.06 for ATG5 cKO; *p*<0.0001. Scale bar, 500μm.

## Discussion

Here, we show that autophagy is necessary for the morphological maintenance and axon guidance of mDA neurons in mice. More importantly, we reveal for the first time that autophagy is required for GC responsiveness to guidance cues to regulate proper axon guidance.

Axon development is a highly complex process that requires tight regulation and cellular homeostasis. It involves axonal elongation, pathfinding and branching. These processes require dynamic changes in the cytoskeleton (Dent and Kalil, 2001) and cytoplasmatic membrane (Pfenninger et al., 2003), mitochondrial energy production homeostasis (Vaarmann et al., 2016) and axonal/GC proteome turnover (Campbell and Holt, 2001), all of which have in common autophagy as a key regulator (Jin et al., 2017; Kast and Dominguez, 2018; Lee et al., 2012). Our observations that autophagy is present in mDA developing neurons and axons and that autophagy markers display a gradual increase during the mDA axonal developmental window, peaking at P7 with a posterior decrease toward adulthood, underscore the relevance of autophagy.

Autophagy dysregulation by *ATG5 or Atg12* knockout in mDA neurons resulted in enlarged axons and GCs, both *in vitro* and *in vivo.* Interestingly, dopaminergic axonal profile enlargements were also observed in *Dat*^*Cre*/+^*ATG7*^*flox/flox*^ animals (Hernandez et al., 2012). These observations underscore the notion that autophagy biogenesis in neurons occurs distally in axons with retrograde transport of autophagosomes to the soma where lysis occurs (Maday et al., 2012). This is particularly relevant for mDA neurons that display long and highly metabolic axons (Pacelli et al., 2015). It has been previously shown that mitophagy occurs locally in the distal axons of mDA neurons (Ashrafi et al., 2014), and it is conceivable that macroautophagy also occurs in the distal growing axon and GCs, where there is the necessity of rapid turnover of cytoskeletal proteins. In line with this, our data show that the GCs of autophagy-deficient cells displayed aberrant increased numbers of microtubule loops, and it has been previously shown that GC area depends on microtubule looping (Purro et al., 2008). We also observed an increased area of these loops. The increased looping prevalence and size can explain the enlargements that we observed, although we cannot exclude the possibility of protein and organelle accumulation due to lack of autophagic degradation.

The lack of autophagy not only affected the GC area but also led to a lower axonal complexity by decreasing the number of branching points. No difference was observed, however, in the initial number of axons that grew out of the explant or in the distance that the axons grew. These data are also in line with our *in vivo* analysis in the developing P11 mice. ATG5 cKO mice displayed a decreased number of axonal profiles in the striatum, corroborating the *in vitro* Sholl analysis. Additionally, using time-lapse imaging of axons derived from cultured explants of ATG5 cKO mice and their littermates, we observed a decreased displacement rate of GCs following autophagy impairment. Altogether, these data strongly argue that autophagy is required for proper mDA axonal growth. As already mentioned, this process requires very dynamic structural plasticity, and our data suggest that such plasticity would be achieved only with the involvement of autophagy. Our data are in contrast with those of Ban et al. (Ban et al., 2013) and Yang et al. (Yang et al., 2017), where autophagy seemed to negatively regulate axon growth in cortical neurons. However, this difference might be explained by neuronal type-specific differences (dopaminergic neurons vs. cortical neurons) and by the experimental approach. Ban et al. made use of pharmacological (rapamycin) and interference RNA approaches, both of which can act on more than one target. Rapamycin is an mTOR inhibitor, which in turn is a protein hub controlling the phosphorylation of many proteins important in cell homeostasis, including, but not limited to, autophagy (Saxton and Sabatini, 2017). Yang et al. identified a microRNA that targets *ATG12*. Nevertheless, microRNAs are known to be polyvalent and are able to regulate more than one target. To minimize off-target effects from our analysis, we used a genetic Cre model, where we specifically knocked out ATG5, a core protein required for autophagy. Hence, ATG5 cKO results in halting autophagy, different from models of autophagy adaptor and/or regulator protein interference that likely partially disrupt autophagy. Mirroring our data with ATG5 cKO are our results with the CRISPR-Cas9 gene editing of ATG12, another core autophagy protein. It should also be mentioned that changes in axonal morphology and growth do not result from the degeneration of mDA neurons in the midbrain following a lack of autophagy. While it is well established that many neurodegenerative disorders are linked to metabolic stress (Jha et al., 2017) and autophagy dysfunction (Tan et al., 2014), the numbers of TH+ neurons in the VTA and SNpc were unaltered. Additionally, autophagy conditional knockout mice models specific to dopaminergic neurons (conditional knockouts of *ATG7* under DAT and TH promoters) display degeneration signs only after 1 month in the striatum and after at least 4 months at the midbrain level (Friedman et al., 2012; Hernandez et al., 2012). Although these data are from *ATG7* conditional knockouts, both ATG7 and ATG5 act at the same autophagy conjugation machinery, and mice deficient for these proteins display similar phenotypes (Nishiyama et al., 2007). Therefore, considering our stereological data and the observations from other studies in which degeneration only occurs later than our analysis age (P11), this suggests that the effects we observe are developmental defects rather than degeneration.

Axon growth and guidance, although conceptually distinct processes, occur jointly, making it difficult to completely separate one from the other. Interestingly, rapid local degradation in the axon by the proteasome system has been implicated in chemotropic responses to cues (Campbell and Holt, 2001). Surprisingly, autophagy has not been addressed in such a paradigm, and the specific involvement of this degradation pathway in axon guidance has not been studied. Therefore, we challenged control cultures and ATG5 cKO cultures with Sema7a (chemorepulsive) and Netrin-1 (chemoattractant) guidance cues. Such cues are of great importance for mDA axon guidance and for the consolidation of the dopaminergic system (Chabrat et al., 2017; Flores, 2011; Van den Heuvel and Pasterkamp, 2008). Sema7a is a chemorepulsive cue that has been recently shown to be required for dorso-ventral mDA axonal organization in the striatum (Chabrat et al., 2017). By contrast, Netrin-1 is mostly known to be a chemoattractant cue that is required for proper guidance of mDA axons toward several targets, including the striatum and PFC (Flores, 2011). Our data reveal that autophagy is dynamically modulated by guidance cues. Both Sema7a and Netrin-1 exert similar effects on axonal autophagy by leading to an enrichment in LC3 dots (autophagosome-related phenotype) in the GCs. Interestingly, axons that were not adjacent to GCs (40 μm away) showed a decrease in the number of LC3 dots in the same mDA neurons. Such an observation suggests that upon guidance cues interaction with their receptors, autophagosomes might be recruited and relocated to the GCs. Further corroborating such hypotheses are our western blot data showing similar LC3-II levels under baseline conditions and following Sema7a and Netrin-1 applications. Hence, it is possible that the total level of autophagy is not altered, but it might be redirected to where it is mostly needed at the moment. In light of these data, we challenged ATG5 cKO cultures with these guidance cues. Strikingly, when explant cultures were incubated with Sema7a and Netrin-1, ATG5 cKO explants failed to react to these cues as revealed by our time-lapse imaging. By contrast, axons from control explants decreased their average displacement when incubated with Sema7a and increased it when incubated with Netrin-1. Altogether, these observations underscore the importance and requirement for autophagy downstream of these guidance cues in mDA axon guidance. In contrast to Netrin-1, which is a diffusible cue, Sema7a is a membrane-associated GPI-anchored protein. Therefore, we also challenged the axonal response of VTA explants growing on alternating stripes of Sema7a. As we previously showed (Chabrat et al., 2017), wild-type cultures grown on control stripes showed no preference, growing randomly, but those cultured on Sema7a-containing carpets displayed clear avoidance of Sema7a stripes, growing preferentially onto control stripes. Remarkably, the effect on Sema7a is completely blunted in explants from ATG5 cKO mice. This finding and our time-lapse imaging indicate that autophagy is required for normal pathfinding behavior in mDA axons in response to guidance cues. Our data become particularly relevant considering that Ambra1 KO (Fimia et al., 2007), mir505-p3 KO (Yang et al., 2017), Alfy/WDFY3 (Bosco et al., 2016) and ULK1/2 whole-brain KO (Wang et al., 2018), all of which are autophagy regulators, result in axonal defects. Importantly, the possibility cannot be excluded that these effects are non-autophagy related, as these are not core autophagy proteins, meaning that autophagy was still occurring in those models and that these proteins are likely related to regulatory functions outside the scope of autophagy. Indeed, Wang et al. claim that the defects seen in their model are noncanonical. Although autophagy levels are not altered in their model, this does not exclude the possibility that autophagy might still be differentially regulated. Localization, for instance, might be altered, or specific autophagy cargo degradation might be deregulated — none of which would likely appear in overall autophagy marker quantifications. By contrast, all of these studies were performed in KO models that are not specific to a distinct cell population. For this reason, it cannot be excluded that these axonal defects could be related to other nonneuronal specific effects. Indeed, if autophagy in glial guideposts is altered, it could also interfere with cellular organization in the brain, which in turn could lead to axonal pathfinding issues. Indeed, one of the phenotypes described by Bosco et al. (Bosco et al., 2016) with the loss of Alfy/WDFY3 is the disruption in the localization of glial guidepost cells.

In sum, increasing evidence around autophagy in brain development shows that autophagy likely has important roles in axon growth and guidance. Our findings reveal autophagy as a key player in the maintenance of mDA axonal morphology and as a prominent mechanism downstream of guidance cue-related effects. Therefore, autophagy appears to be a central mechanism that tightly regulates mDA system development.

## Materials and Methods

### Animals

All animal experiments were performed in accordance with the Canadian Guide for the Care and Use of Laboratory Animals and were approved by the Université Laval Animal Protection Committee. *Atg5*^*f/f*^ (Hara et al., 2006) and *Dat*^*Cre*/+^*(Zhuang et al., 2005)* mice were genotyped as previously described. *Atg5* cKO mice were generated by intercrossing *Dat*^*Cre*/+^ males *and Atg5f*^*/f*^ females. *Atg5*^*f/f*^ mice were used as controls, and *Dat*^*Cre*/+^*Atg5*^*f/f*^ – *Atg5* cKO mice were used as the experimental animals. *Atg5* transgenic mice were identified by PCR with forward primers in the *Atg5* sequence, 5-GAATATGAAGGCACACCCCTGAAATG-3, and reverse primers 5-ACAACGTCGAGCACAGCTGCGCAAGG-3 and 5-GTACTGCATAATGGTTTAACTCTTGC-3, identifying heterozygous cassettes and those homozygous for *floxed*, respectively.

### Tissue analysis

Mouse brains at P1 were incubated in 4% paraformaldehyde in PBS at 4 °C, followed by cryoprotection in 30% sucrose in PBS, before they were frozen in dry ice. For mice older than P1, perfusion using 4% paraformaldehyde in PBS was instead performed. After cryostat sectioning at 60 μm, sections were washed in PBS and then blocked with 1% normal donkey serum (NDS) and 0.2% Triton X-100 for at least 30 min. When rabbit anti-LC3 was used, sections were instead blocked with 1% normal donkey serum (NDS), 5% milk and 0.2% Triton X-100 for at least 30 min. Primary incubation was performed overnight at 4 °C, except for rabbit anti-LC3, for which a two overnight incubation at 4 °C was performed. The primary antibodies used in this study were rabbit anti-TH (Pel-Freez Biologicals, P40101, 1:1000), sheep anti-TH (Pel-Freez Biologicals, P60101, 1:1000) and rabbit anti-LC3 (Abgent, AP1801a, 1:250). The secondary antibodies used in this study were donkey Alexa-Fluor-488, donkey Alexa-Fluor-555 or donkey Alexa-Fluor-647 (Life Technologies), used at 1:400, and donkey Cy3 and donkey FITC (Jackson Immunoresearch), used at 1:200.

### Stereological neuron counting

The number of TH+ and GFP+ neurons within the VTA and SNc of mutants and controls was quantified using the optical fractionator stereological method (Stereo Investigator; MBF Bioscience). The following analytic parameters were applied: 100×100 μm counting frame size, 15 μm optical dissector height, and 1 in 3 section interval. The coefficients of error (Gundersen m=1) were less than 0.10.

### Western blotting

Ventral midbrain from control *Pitx3*^*GFP*/+^ mice and from *Dat*^+/+^ *Atg5*^*f/f*^ and *Dat*^*Cre*/+^*Atg5*^*f/f*^ brains at E12, E14, E18, P1, and P7 and adult mice were dissected, and samples were then snap frozen. Sample lysis was performed in radioimmunoprecipitation (RIPA) buffer complemented with protease inhibitor and phosphatase inhibitor cocktails (Roche) (50 mM Tris-HCl, 150 mM NaCl, 1% NP-40, 0.5% sodium deoxycholate, 0.1% SDS, 2 mM EDTA, 50 mM NaCl, pH 7.4). Quantification of protein content in the samples was performed using a DC-protein assay (Bio-Rad). Protein extracts (15 μg or 30 μg for LC3-II detection) were separated by SDS-polyacrylamide gel electrophoresis (12% and 15% SDS-PAGE Tris-glycine gels). Nitrocellulose and PVDF (for LC3-II detection) membranes (Bio-Rad) were used for the transfer. Blots were blocked in 7% milk in 1× TBS and 0.01% Tween-20 (Sigma) and immunostained overnight at 4 °C with primary antibodies in blocking solution. For LC3-II detection, 2 overnight incubation was performed. The primary antibodies mouse anti-actin (1:10,000; Millipore, MAB1501), rabbit anti-LC3 (1:500; Abcam, ab AP1801a), sheep anti-TH (1:1000; Pel-Freez Biologicals, P60101), rabbit anti-ATG5 (1:500; Abcam, AP1812b) and rabbit anti-p62 (1:1000; Proteintech, 18420-1-AP) were diluted in blocking solution. Blots were washed 3 × 5 min with 1× TBS and 0.01% Tween-20, and immune complexes were detected with species-appropriate secondary antibodies conjugated to HRP, including goat anti-rabbit HRP (1:3000; CST 7074), goat anti-mouse HRP (1:5000; Life Technologies, G-21040) and donkey anti-sheep HRP (1:5000; Santa Cruz, sc-2473). Membranes were covered with ECL for 1 min (Western Lightning Plus-ECL, PerkinElmer), and chemiluminescence was then documented by exposing the membranes to Pierce CL-Xposure films (Thermo Scientific). Films were scanned and analyzed using the ImageJ64 program.

### Ventral midbrain primary cell and explant cultures

P1 mouse ventral mesencephalons were dissected in L-15 medium (Life Technologies) and dissociated in papain solution [12 U/mL papain (Worthington Biochemical), 250 U/mL DNase I type IV (Sigma), 3.5 mM L-cysteine (Sigma-Aldrich), 0.215% NaHCO_3_ (Sigma-Aldrich), 5 mM EDTA (Life Technologies), 0.2% Phenol Red (Sigma-Aldrich), 1 mM sodium pyruvate (Life Technologies), 1.8 mg/mL D-glucose (Sigma-Aldrich), 50 U/mL penicillin and 50 μg/mL streptomycin (Life Technologies) in HBSS without Ca^2+^ and Mg^2+^ (Life Technologies)]. Mechanical trituration was then performed in trituration solution [0.2% BSA (Sigma-Aldrich), 50 U/mL penicillin, 50 μg/mL streptomycin, 1 mM sodium pyruvate, 1.8 mg/mL glucose, and 250 U/mL DNase I type IV in neurobasal medium (Life Technologies)]. Cells purification was performed using BSA columns [1.8% (wt/vol) BSA, 50 U/mL penicillin, 50 μg/mL streptomycin, 1 mM sodium pyruvate, 1.8 mg/mL glucose, 250 U/mL DNase I type IV, 3 mM NaOH (Fisher Scientific) in neurobasal medium] and centrifugation at 800 × *g* for 5 min. Cells were seeded on top of 12-mm coverslips coated with 0.003% poly-L-ornithine (Sigma) and 10 μg/mL laminin (Life Technologies) (Fisher Scientific) and maintained at 37 °C in a humidified atmosphere of 5% CO_2_ in complete growth medium [10% (vol/vol) FBS (Life Technologies), 1:50 B27 supplement (Life Technologies), 1:100 GlutaMAX (Life Technologies), and1.2 mg/mL D-glucose in neurobasal medium] at low density (3,000,000 cells/mL). After 3DIV cells were fixed for 30 min at 4 °C in fixative solution (4% PFA, 4% sucrose, in 1× PBS), immunostaining was performed. TH (sheep anti-TH; Pel-Freez Biologicals, P60101; 1:1000) and LC3 (rabbit anti-LC3; Abgent, AP1801a; 1:500) colabeling was performed with two overnight incubations in 5% milk, 1% NDS and 0.2% Triton X-100 in 1× PBS. TH (sheep anti-TH; Pel-Freez Biologicals, P60101; 1:1000) and tubulin (mouse anti-alpha-tubulin, Sigma-Aldrich, T5168; 1:250) colabeling was performed overnight with 1% NDS and 0.2% Triton X-100 in 1× PBS. LC3 dots and tubulin loops were manually counted using confocal images and ImageJ software.

Embryonic ventral midbrain explants were dissected from E14.5 Pitx3^*GFP*/+^ (electroporation assay), *Dat*^*Cre*/+^ *Atg5*^*f/f*^ and *Dat*^+/+^ *Atg5*^*f/f*^ embryos in ice-cold L15 with 5% FBS. Explants were grown on 12-mm diameter glass coverslips coated with 20 μg/mL laminin for 2D cultures used for time-lapse imaging or coated with 7 μl of Matrigel™ (BD Biosciences, Mississauga, ON, Canada) for 3D cultures used for morphological analysis. 2D explants were then cultured in 1 ml of neurobasal medium complemented with B27, PenStrep, GlutaMAX, sodium pyruvate, and FBS (0.4% methyl cellulose 1500 centipoise, 86.8% neurobasal medium, 5% PenStrep, 2% B27, 0.2% GlutaMAX, 1% sodium pyruvate, and 5% FBS) for 1 day at 37 °C, 5% CO_2_ prior to time-lapse imaging. 3D explants were each covered by 7 μl of Matrigel and cultured in 1 ml of neurobasal medium complemented with B27, PenStrep, GlutaMAX, sodium pyruvate, and FBS (86.8% neurobasal medium, 5% P/ S, 2% B27, 0.2% GlutaMAX, 1% sodium pyruvate, and 5% FBS) for 3 days at 37 °C, 5% CO_2_. After 3 DIVs, 3D explants were fixed for 30 min at 4 °C in fixative solution (4% PFA, 4% sucrose, in 1× PBS). TH was immunostained by overnight incubation with sheep anti-TH (Pel-Freez, 1:1000) in 1% NDS and 0.2% Triton X-100 in 1× PBS. Time-lapse analysis was performed on DiC images using MTrackJ plugin. Sholl analysis was performed on confocal images of the TH signal using the Neurite-J plug-in.

### Stripe assay

The stripe assay was performed as described by Knoll, B. et al. (Knöll et al., 2007). Glass coverslips were coated with alternating stripes (IgG or Sema7a 100 μg/ml, R&D Systems), which were then covered by laminin (20 μg/ml). Explants from ATG5 cKO mice and control littermates were seeded on top of the coatings, cultured for 3 days, and fixed with 4% PFA, 4% sucrose, in 1× PBS before immunostaining. TH was immunostained by overnight incubation with sheep anti-TH (Pel-Freez, 1:1000) in 1% NDS and 0.2% Triton X-100 in 1× PBS. For quantification, the number of neurites terminating on control vs. Sema7a stripes was manually counted for each explant using confocal images and ImageJ software.

### Statistical analyses

GraphPad Prism 7.0 (GraphPad Software, La Jolla, CA, USA) software was used for the statistical analyses. The differences between two groups were determined by Student’s t test. Sholl analysis and time-lapse data were analyzed by two-way analysis of variance, and the Sidak test was used for post hoc comparisons. All data are represented as mean ± SEM, and significance is defined as *p < 0.05, **p < 0.01, or ***p < 0.001.

### Microscopes

Immunofluorescence images were acquired using a Zeiss LSM5 Pascal confocal microscope or Zeiss LSM700 confocal microscope and then processed using ImageJ software and Adobe Photoshop CS6. Bright-field pictures were acquired using a Leica DMRB equipped with a digital camera. Bright-field images were acquired using a TISSUEscope™ 4000 (Huron Technologies).

## Acknowledgments

We thank Dr. Marina Snapyan for the design and cloning of gRNAs, and the members of AS and ML labs for helpful comments. We also thank Veronique Rioux, for her technical assistance. This work was supported by the Natural Sciences and Engineering Research Council of Canada (RGPIN-2018-06262 to M.L.) and the Canadian Institute of Health Research (31120 to M.L. and PJT-153026 to A.S.). MSP was partially supported by Pierre Durand and CTRN fellowships from the Faculty of Medicine of Université Laval. ML is a career awardee of the FRQS in partnership with Parkinson Québec (34974). The authors declare no competing interest.

## Author Contributions

Conceptualization, M.S.P., A.S. and M.L.; investigation, M.S.P., A.S. and M.L.; writing of the manuscript, M.S.P., A.S. and M.L.; supervision, A.S. and M.L.

## Supplementary Figures

**Fig. S1.**
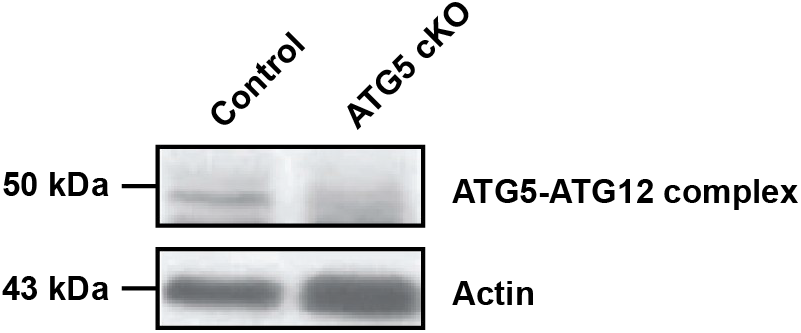
Cre recombinase under Dat transporter promoter effectively knockouts floxed *Atg5* gene. Western blot of autophagic marker ATG5-ATG12 complex displays lack of ATG5-ATG12 protein complex in P7 ATG5 cKO animals in comparison to control littermates (n= 3 animals per group).

**Fig. S2.**
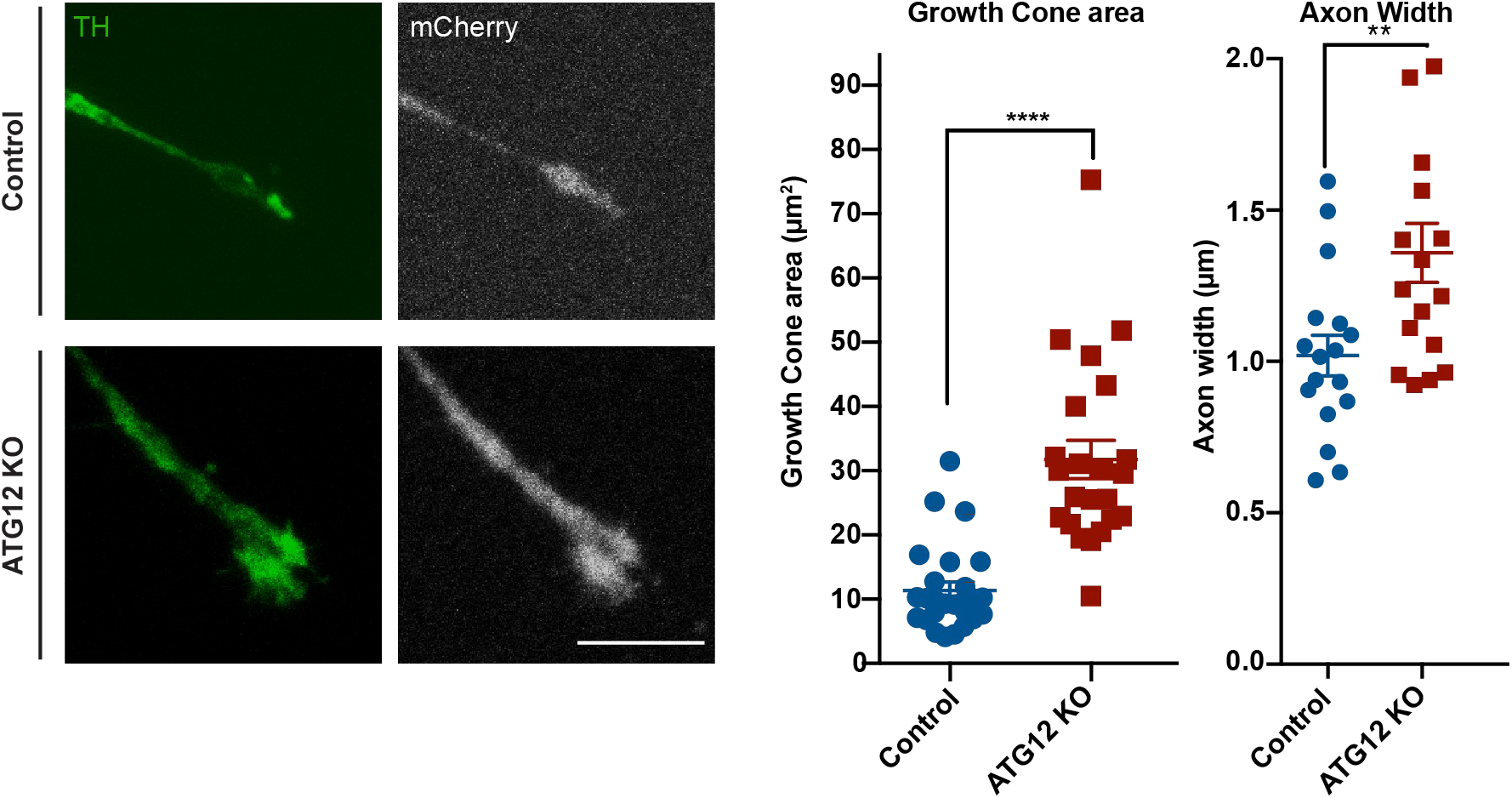
Autophagy ablation by CRSPR/Cas9 *Atg12* knockout in mDA explants leads to aberrant axon morphology. E14 VTA from control mice were dissected into explants, electroporated with expression plasmids containing Cas9 and *LacZ* guide as a control or *Atg12* guides for experimental explants and cultured in 3D collagen matrix for 3DIV prior to fixation and ICC for TH labelling. mCherry signal is the result of the direct overexpression of the plasmid marker without ICC. GC area: n=26 GCs for control samples and n=23 GCs for ATG12 KO samples from an average of 4 explants from 3 independent experiments (mean values= 11.37 for control and 31.73 for ATG12 KO; p<0.0001; two-tailed t test). Axon width: n=17 axons for control samples and n=17 axons for ATG12 KO samples from 4 explants from 3 independent experiments (mean values= 1.01 for control and 1.35 for ATG12 KO; p= 0.0071; two-tailed t-test). Scale bar, 10 μm.

## References

Ashrafi, G., Schlehe, J.S., LaVoie, M.J., and Schwarz, T.L. (2014). Mitophagy of damaged mitochondria occurs locally in distal neuronal axons and requires PINK1 and Parkin. J. Cell Biol. 206, 655–670.

Ban, B.-K., Jun, M.-H., Ryu, H.-H., Jang, D.-J., Ahmad, S.T., and Lee, J.-A. (2013). Autophagy negatively regulates early axon growth in cortical neurons. Mol. Cell. Biol. 33, 3907–3919.

Bosco, J.R., Hirose-Ikeda, M., Mita, T., Simonsen, A., Fox, L.M., Yoon, M.S., Ichimura, Y., Lystad, A.H., Mason, C.A., Komatsu, M., et al. (2016). Autophagy linked FYVE (Alfy/WDFY3) is required for establishing neuronal connectivity in the mammalian brain. Elife 5, 1–25.

Campbell, D.S., and Holt, C.E. (2001). Chemotropic Responses of Retinal Growth Cones Mediated by Rapid Local Protein Synthesis and Degradation. 32, 1013–1026.

Chabrat, A., Brisson, G., Doucet-Beaupré, H., Salesse, C., Schaan Profes, M., Dovonou, A., Akitegetse, C., Charest, J., Lemstra, S., Côté, D., et al. (2017). Transcriptional repression of Plxnc1 by Lmx1a and Lmx1b directs topographic dopaminergic circuit formation. Nat. Commun. 8.

Dent, E.W., and Kalil, K. (2001). Axon Branching Requires Interactions between Dynamic Microtubules and Actin Filaments. 21, 9757–9769.

Dent, E.W., Callaway, J.L., Szebenyi, G., Baas, P.W., Kalil, K., Dent, E.W., and Callaway, J.L. (1999). Reorganization and Movement of Microtubules in Axonal Growth Cones and Developing Interstitial Branches. J. Neurosci. 19, 8894–8908.

Fimia, G.M., Stoykova, A., Romagnoli, A., Giunta, L., Di Bartolomeo, S., Nardacci, R., Corazzari, M., Fuoco, C., Ucar, A., Schwartz, P., et al. (2007). Ambra1 regulates autophagy and development of the nervous system. Nature 447, 1121–1125.

Flores, C. (2011). Role of netrin-1 in the organization and function of the mesocorticolimbic dopamine system. J. Psychiatry Neurosci. 36, 296–310.

Friedman, L.G., Lachenmayer, M.L., Wang, J., He, L., Poulose, S.M., Komatsu, M., Holstein, G.R., and Yue, Z. (2012). Disrupted autophagy leads to dopaminergic axon and dendrite degeneration and promotes presynaptic accumulation of α-synuclein and LRRK2 in the brain. J. Neurosci. 32, 7585–7593.

Hara, T., Nakamura, K., Matsui, M., Yamamoto, A., Nakahara, Y., Suzuki-Migishima, R., Yokoyama, M., Mishima, K., Saito, I., Okano, H., et al. (2006). Suppression of basal autophagy in neural cells causes neurodegenerative disease in mice. Nature 441, 885–889.

He, M., Ding, Y., Chu, C., Tang, J., Xiao, Q., and Luo, Z.-G. (2016). Autophagy induction stabilizes microtubules and promotes axon regeneration after spinal cord injury. Proc. Natl. Acad. Sci. 201611282.

Hernandez, D., Torres, C. a, Setlik, W., Cebrián, C., Mosharov, E. V, Tang, G., Cheng, H.-C., Kholodilov, N., Yarygina, O., Burke, R.E., et al. (2012). Regulation of presynaptic neurotransmission by macroautophagy. Neuron 74, 277–284.

Van den Heuvel, D.M. a, and Pasterkamp, R.J. (2008). Getting connected in the dopamine system. Prog. Neurobiol. 85, 75–93.

Hu, Z., Cooper, M., Crockett, D.P., and Zhou, R. (2004). Differentiation of the midbrain dopaminergic pathways during mouse development. J. Comp. Neurol. 476, 301–311.

Jha, S.K., Jha, N.K., Kumar, D., Ambasta, R.K., and Kumar, P. (2017). Linking mitochondrial dysfunction, metabolic syndrome and stress signaling in Neurodegeneration. Biochim. Biophys. Acta - Mol. Basis Dis. 1863, 1132–1146.

Jin, E.J., Kiral, F.R., and Hiesinger, P.R. (2017). The Where , What , and When of Membrane Protein Degradation in Neurons.

Kast, D.J., and Dominguez, R. (2018). The Cytoskeleton-Autophagy Connection. 27.

Kaur, J., and Debnath, J. (2015). Autophagy at the crossroads of catabolism and anabolism. Nat. Rev. Mol. Cell Biol.

Khan, M.M., Strack, S., Wild, F., Hanashima, A., Gasch, A., Brohm, K., Reischl, M., Carnio, S., Labeit, D., Sandri, M., et al. (2014). Role of autophagy, SQSTM1, SH3GLB1, and TRIM63 in the turnover of nicotinic acetylcholine receptors. Autophagy 10, 123–136.

Klionsky, D.J., Abdelmohsen, K., Abe, A., Abedin, M.J., Abeliovich, H., Arozena, A.A., Adachi, H., Adams, C.M., Adams, P.D., and Adeli, K., et al. (2016). Guidelines for the use and interpretation of assays for monitoring autophagy (3rd edition). Autophagy 12, 1–222.

Knöll, B., Weinl, C., Nordheim, A., and Bonhoeffer, F. (2007). Stripe assay to examine axonal guidance and cell migration. Nat. Protoc. 2, 1216–1224.

Lamb, C.A., Dooley, H.C., and Tooze, S.A. (2013). Endocytosis and autophagy: Shared machinery for degradation. BioEssays 35, 34–45.

Lee, J., Giordano, S., and Zhang, J. (2012). Autophagy, mitochondria and oxidative stress: cross-talk and redox signalling. Biochem. J 540, 523–540.

Maday, S., Wallace, K.E., and Holzbaur, E.L.F. (2012). Autophagosomes initiate distally and mature during transport toward the cell soma in primary neurons. J. Cell Biol. 196, 407–417.

Mizushima, N., and Levine, B. (2010). Autophagy in mammalian development and differentiation. Nat. Cell Biol. 12, 823–830.

Nishiyama, J., Miura, E., Mizushima, N., Watanabe, M., and Yuzaki, M. (2007). Aberrant membranes and double-membrane structures accumulate in the axons of Atg5-null Purkinje cells before neuronal death. Autophagy 3, 591–596.

Pacelli, C., Gigue, N., and Slack, R.S. (2015). Elevated Mitochondrial Bioenergetics and Axonal Arborization Size Are Key Contributors to the Vulnerability of Dopamine Neurons Article Elevated Mitochondrial Bioenergetics and Axonal Arborization Size Are Key Contributors to the Vulnerability of Dopamine. 2349–2360.

Pfenninger, K.H., Laurino, L., Peretti, D., Wang, X., Rosso, S., Morfini, G., Cáceres, A., and Quiroga, S. (2003). Regulation of membrane expansion at the nerve growth cone.

Prestoz, L., Jaber, M., and Gaillard, A. (2012). Dopaminergic axon guidance: which makes what? Front. Cell. Neurosci. 6, 32.

Purro, S.A., Ciani, L., Hoyos-Flight, M., Stamatakou, E., Siomou, E., and Salinas, P.C. (2008). Wnt Regulates Axon Behavior through Changes in Microtubule Growth Directionality: A New Role for Adenomatous Polyposis Coli. J. Neurosci. 28, 8644–8654.

Saxton, R.A., and Sabatini, D.M. (2017). mTOR Signaling in Growth, Metabolism, and Disease Robert. Cell 168, 960–976.

Sutherland, D.J., Pujic, Z., and Goodhill, G.J. (2014). Calcium signaling in axon guidance. Trends Neurosci. 37, 424–432.

Tamariz, E., and Varela-Echavarría, A. (2015). The discovery of the growth cone and its influence on the study of axon guidance. Front. Neuroanat. 9, 1–9.

Tan, C.C., Yu, J.T., Tan, M.S., Jiang, T., Zhu, X.C., and Tan, L. (2014). Autophagy in aging and neurodegenerative diseases: Implications for pathogenesis and therapy. Neurobiol. Aging 35, 941–957.

Tojima, T., Hines, J.H., Henley, J.R., and Kamiguchi, H. (2011). Second messengers and membrane trafficking direct and organize growth cone steering. Nat. Rev. Neurosci. 12, 191–203.

Vaarmann, A., Mandel, M., Zeb, A., Wareski, P., Liiv, J., Kuum, M., Antsov, E., Liiv, M., Cagalinec, M., Choubey, V., et al. (2016). Mitochondrial biogenesis is required for axonal growth. 1981–1992.

Wang, B., Iyengar, R., Li-harms, X., Hyuck, J., Wright, C., Lavado, A., and Horner, L. (2018). The autophagy-inducing kinases , ULK1 and ULK2 , regulate axon guidance in the developing mouse forebrain via a noncanonical pathway. 14, 796–811.

Yang, K., Yu, B., Cheng, C., Cheng, T., Yuan, B., Li, K., Xiao, J., Qiu, Z., and Zhou, Y. (2017). Mir505–3p regulates axonal development via inhibiting the autophagy pathway by targeting Atg12. Autophagy 13, 1679–1696.

Zhuang, X., Masson, J., Gingrich, J.A., Rayport, S., and Hen, R. (2005). Targeted gene expression in dopamine and serotonin neurons of the mouse brain. J. Neurosci. Methods 143, 27–32.

